# Transcriptome analysis of sevoflurane exposure effects at the different brain regions

**DOI:** 10.1101/2020.07.15.204040

**Authors:** Hiroto Yamamoto, Yutaro Uchida, Tomoki Chiba, Ryota Kurimoto, Takahide Matsushima, Chihiro Ishikawa, Haiyan Li, Takashi Shiga, Masafumi Muratani, Tokujiro Uchida, Hiroshi Asahara

## Abstract

**Backgrounds:** Sevoflurane is a most frequently used volatile anaesthetics, but its molecular mechanisms of action remain unclear. We hypothesized that specific genes play regulatory roles in whole brain exposed to sevoflurane. Thus, we aimed to evaluate the effects of sevoflurane inhalation and identify potential regulatory genes by RNA-seq analysis.

**Methods:** Eight-week old mice were exposed to sevoflurane. RNA from four medial prefrontal cortex, striatum, hypothalamus, and hippocampus were analysed using RNA-seq. Differently expressed genes were extracted. Their gene ontology terms and the transcriptome array data of the cerebral cortex of sleeping mice were analysed using Metascape, and the gene expression patterns were compared. Finally, the activities of transcription factors were evaluated using a weighted parametric gene set analysis (wPGSA). JASPAR was used to confirm the existence of binding motifs in the upstream sequences of the differently expressed genes.

**Results:** The gene ontology term enrichment analysis result suggests that sevoflurane inhalation upregulated angiogenesis and downregulated neural differentiation in the whole brain. The comparison with the brains of sleeping mice showed that the gene expression changes were specific to anaesthetized mice. Sevoflurane induced *Klf4* upregulation in the whole brain. The transcriptional analysis result suggests that KLF4 is a potential transcriptional regulator of angiogenesis and neural development.

**Conclusions:** *Klf4* was upregulated by sevoflurane inhalation in whole brain. KLF4 might promote angiogenesis and cause the appearance of undifferentiated neural cells by transcriptional regulation. The roles of KLF4 might be key to elucidating the mechanisms of sevoflurane induced functional modification in the brain.

## Introduction

Sevoflurane is a most frequently used volatile anaesthetics in general anaesthesia. Some reports discussed the peri-operative adverse effects of sevoflurane such as post-operative delirium and cognitive disorders, although whether anesthetics themselves cause peri-operative adverse effects is still controversial [1–3]. Several membrane receptors such as the γ-aminobutyric acid type A receptor, nicotinic AchR, hyperpolarization-activated cyclic nucleotide-gated channels have been reported to be potential targets of sevoflurane [4–9]. However, receptor-based molecular mechanisms have not sufficiently explained these phenomena. Furthermore, although many reports have evaluated effects of sevoflurane, the analyses were focused only on limited parts of the brain; hence, comparison of the effects of sevoflurane between multiple anatomical sites at the same time is difficult.

In this study, we focused on a simultaneous analysis of the effects of sevoflurane on the gene expression changes in multiple anatomical sites of the brain. The medial prefrontal cortex (MPFC), hippocampus, striatum, and hypothalamus were chosen as targets of the analysis, as these parts were frequently used for evaluating the effects of volatile anaesthetics [10–13]. We hypothesized that some common genes play important roles in whole brain exposed to sevoflurane and are potential regulators of the sevoflurane-induced changes in brain function. Moreover, we compared gene expression patterns between sevoflurane-induced anaesthesia and physiological sleeping to identify the specific genes that showed expression changes in brains exposed to sevoflurane.

Therefore, in this study, we performed a genome-wide transcriptional analysis of murine brains tissues samples from four parts of the brain of mice that inhaled sevoflurane. Differently expressed genes (DEGs) and enriched gene groups were compared between the four parts of the brain. We applied the same analysis to the transcriptome array data of sleeping mice, and identified specific gene expression changes in brains exposed to sevoflurane. Finally, we evaluated the effects of the transcription factors on their target genes, and confirmed the existence of consensus-binding motifs in the upstream sequences of DEGs. Herein, we report our success in identifying KLF4 as a specific responsible transcription factor that potentially promotes angiogenesis and induces the appearance of undifferentiated neural cells.

## Materials and Methods

### Approval for the animal experiments

All the animal experiments in this study were conducted in accordance with the Guidelines for Proper Conduct of Animal Experiments (Science Council of Japan) and approved by the Center for Experimental Animals of Tokyo Medical and Dental University. (Approval No.A2017-131A)

### Experimental conditions and preparation of brain samples

Eight-week old mice (C57BL/6J) were purchased from Sankyo Labo (Tokyo, Japan). Six mice were assigned into two groups, the control (n=3) and sevoflurane inhalation groups (n=3). For the sevoflurane group, the mice were put in a box with 2.5% sevoflurane / 40% oxygen for 3 hours. The body temperature was measured and sustained within the range of ±0.5°C by using a body warming machine. For the control group, the mice were put in a box with normal air and stayed in the box without food or water for 3 hours. After the treatments, all the mice were immediately killed through cervical dislocation, and their whole brains were removed. The brain samples were cut into 2 mm slices, and the medial prefrontal cortex, striatum, hippocampus, and hypothalamus were punched out, referring to the methods of Ishikawa et al [14].

### RNA extraction from brain tissue sections and RNAseq analysis

RNA was extracted from brain tissue sections by using TRIZOL (ThermoFisher, Waltham, MA, USA) and 500ng total RNA was used for the subsequent preparation. RNA-seq libraries were prepared with a rRNA-depletion kit (E6310, New England Biolabs Japan, Tokyo, Japan) and a directional library synthesis kit (E6310, New England Biolabs Japan). The RNA libraries were sequenced using NextSeq500 High-output kit v2 for 2 × 36 base reads.

### Mapping FASTQ data and calculating gene expressions

The adapters in the FASTQ files were trimmed using the TrimGalore software (https://www.bioinformatics.babraham.ac.uk/projects/trim_galore/). The FASTQ files were mapped to mouse genomes (mm10) by using the STAR software [15] (https://github.com/alexdobin/STAR), and the amount of each transcript was calculated with the RSEM software [16] (https://github.com/deweylab/RSEM).

### Extracting DEGs on iDEP.91

The counted data were transformed with EdgeR (log_2_ [counts per million (CPM) + 4]), and principal components analysis (PCA) plots were depicted. DEGs were extracted using DESeq2. All these steps were performed with iDEP.91 [17] (http://bioinformatics.sdstate.edu/idep/). Venn-diagrams were used to depict the upregulated and downregulated DEGs.

### Sequencing data

The raw sequencing data were submitted to the DNA Data Bank Japan (DDBJ; http://www.ddbj.nig.ac.jp) under accession No.DRA010292.

### Gene ontology term enrich analysis using Metascape

The extracted DEGs were analysed with Metascape[18] (http://metascape.org/gp/index.html#/main/step1). A gene ontology (GO) term enrichment analysis was performed, and a Circos plot was drawn.

### Extraction of DEGs from sleeping mice

The transcriptome array data of the unbound fractions of immunoprecipitation for the cerebral cortices of waking or sleeping mice (GSE69079) were used in the analysis [19]. The expression data were normalized, and DEGs were selected using DESeq2 in iDEP. 91. Venn-diagrams were used to depict the DEGs of the MPFCs of anesthetized mice and cerebral cortices of the sleeping mice.

### Weighted parametric gene set analysis (wPGSA) analysis of DEGs in the brains of mice that inhaled sevoflurane

For the fold changes data, the expression changes of the target genes of each transcription factor were calculated and its activity (T-score) was estimated using a weighted parametric gene set analysis (wPGSA[20]: http://wpgsa.org/). Expressed transcription factors in the brain were extracted from the analysed data. The transcription factors with T-scores of > 2.0 and false discovery rates (FDRs) of < 0.1 were regarded as active transcription factors, while those with T-scores of < −2.0 and FDR of < 0.1 were regarded as suppressive transcription factors. Moreover, the transcription factors included in the DEGs were extracted from active and suppressive transcription factors and, regarded as responsible for anaesthetic effects.

### Detection of the consensus-binding motifs of Klf4 in the upstream sequences of DEGs

The consensus-binding motifs of Klf4 were referred from JASPAR (http://jaspar.genereg.net/). The 1000-bp upstream sequences of the DEGs annotated with the GO terms “angiogenesis” and “head development” were analysed using JASPAR and the existence of Klf4 binding motifs was confirmed. We regarded the motifs with scores of > 8 as candidate binding motifs for Klf4.

### Statistical Analyses

In extracting DEGs from RNA-seq data and differently activating transcription factors from the wPGSA analysed data, we considered FDRs of < 0.1 as statistically significant.

## Results

### Genome-wide transcriptome analysis for the brains of mice that inhaled sevoflurane

To investigate the sevoflurane-induced gene expression changes in the brain, three 8 week old male mice that inhaled sevoflurane for 3 hours or the control mice were killed, and their brains were removed. The brain tissue samples were cut into 2 mm slices, and four parts of the brain (hippocampus, hypothalamus, medial prefrontal cortex and striatum) were punched out for RNA extraction. We performed a genome-wide transcriptional analysis with next generation sequencing, and confirmed the proper RNA extraction from each brain area by the PCA plot (Fig. 1A, S1 FigA and B). DEGs were extracted on the basis of the criteria of FDR < 0.1(S1 Table). Among the upregulated DEGs, 100, 109, 33, and 314 were expressed in the striatum, MPFC, hypothalamus and hippocampus, respectively. Among the downregulated DEGs, 93, 121, 18, and 502 were expressed in the striatum, MPFC, hypothalamus, hippocampus, respectively (Fig. 1B, C, D, and E, S2 Table). The highest number of DEGs was found in the hippocampus; and the lowest number in the hypothalamus. These results suggest that the gene expression in the hippocampus was the most-influenced and that in the hypothalamus was the least-influenced by sevoflurane inhalation.

**Fig1.**
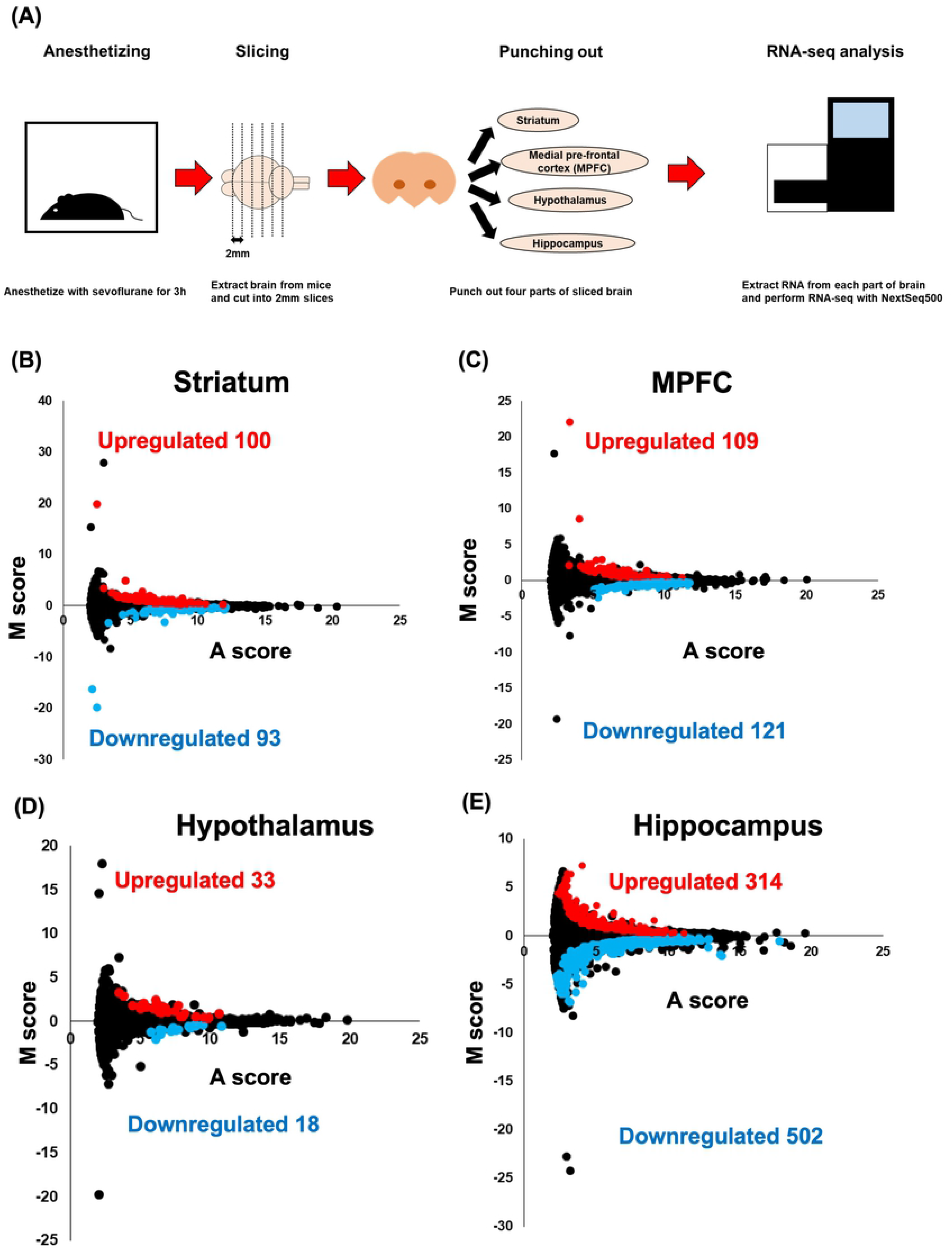
RNA-seq analysis for brains exposed to sevoflurane. (A)The work flow of the RNA-seq analysis for the anesthetized mice. After anaesthetizing with 2.5% of sevoflurane and 40% oxygen for 3 hours, the brains were removed and sliced into 2 mm pieces. The striatum, medial prefrontal cortex (MPFC), hypothalamus and hippocampus were punched out. RNA was extracted from the punched out samples and RNA-seq was performed using NextSeq500. (B)~(E) FASTQ files were mapped using STAR, differently expressed genes (DEGs) were extracted using iDEP.91 and MA-plots were drawn for the striatum (B), medial prefrontal cortex (C), hypothalamus (D), and hippocampus (E).

To compare the upregulated DEGs in the different parts of the brain, a Venn-diagram was drawn (Fig. 2A). Thirteen common upregulated genes found in all parts of the brain are shown in Fig. 2B. Sevoflurane inhalation upregulated *Klf4* and *Ne*s in the whole brain. The expression levels of *Klf4* and *Nes* were >2.5 times higher than those in control mice. KLF4 is a famous transcription factor for sustaining the undifferentiated state of iPS cells, known as the “Yamanaka factor”. NES is a protein marker of neural stem cells and rarely expressed in differentiated neural cells [21–23]. The upregulation of these genes suggest the possibility of induction of the appearance of undifferentiated neural cells by sevoflurane [21–24]. Furthermore, to investigate the differences of upregulated DEGs between the different parts of the brain, a GO term enrichment analysis was performed using Metascape[18]. The Circos plot drawn using Metascape showed similarities in the upregulation patterns of the gene expressions in the four parts of the brain (Fig. 2C). As shown in the heatmap, sevoflurane inhalation caused the upregulation of genes annotated as “angiogenesis” and “response to wounding” in all parts of brain (Fig. 2D). The transcription factors KLF4 and KLF2, as well as EDN1, CCN1, and ADAMTS1, were annotated to the GO terms “angiogenesis” and “response to wounding” (S3 Table). These results suggest that sevoflurane inhalation induced the acceleration of angiogenesis via KLF4 and KLF2 in whole brain.

**Figure2.**
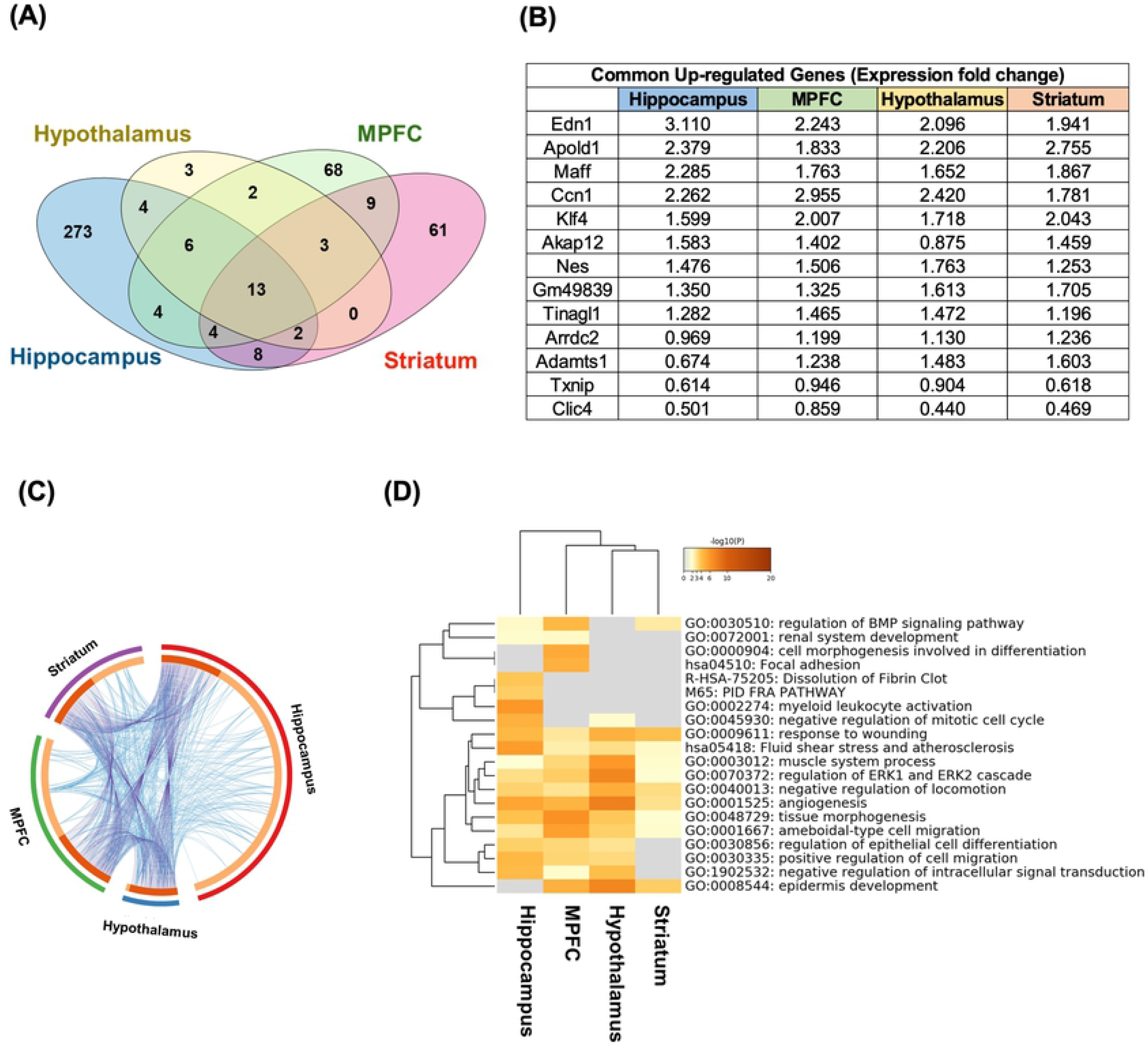
Analysis of the upregulated differently expressed genes (DEGs) in each part of the brain. (A) Venn-diagram for the upregulated differently expressed genes (DEGs) in each part of brain. (B) The 13 genes commonly upregulated in the four parts of the brain and the fold changes (log2) for each gene. (C) Circos plot for the Metascape analysis of upregulated DEGs. The purple line links the same gene that is shared by multiple gene lists. The blue lines link the different genes where they fall into the same ontology term. (D) Heatmap for the gene ontology term analysis of the upregulated DEGs.

Next, downregulated DEGs were compared between the four parts of the brain, and a Venn-diagram was drawn (Fig. 3A). The common downregulated DEGs among whole brain was only the Banp gene (Fig. 3B). Furthermore, to compare the downregulated DEGs between the four parts of the brain, a GO term enrichment analysis was performed with Metascape. As shown in the Circos plot, enriched GO terms were similar among the different parts of the brain (Fig. 3C). Moreover, the heatmap showed that sevoflurane inhalation downregulated the genes annotated as “head development” in whole brain, and those annotated as “axon development” or “synapse organization” in several parts (Fig. 3D and S4 Table). These results indicate that sevoflurane inhalation might suppress neural development, and these effects might reflect the appearance of undifferentiated neural cells.

**Figure3.**
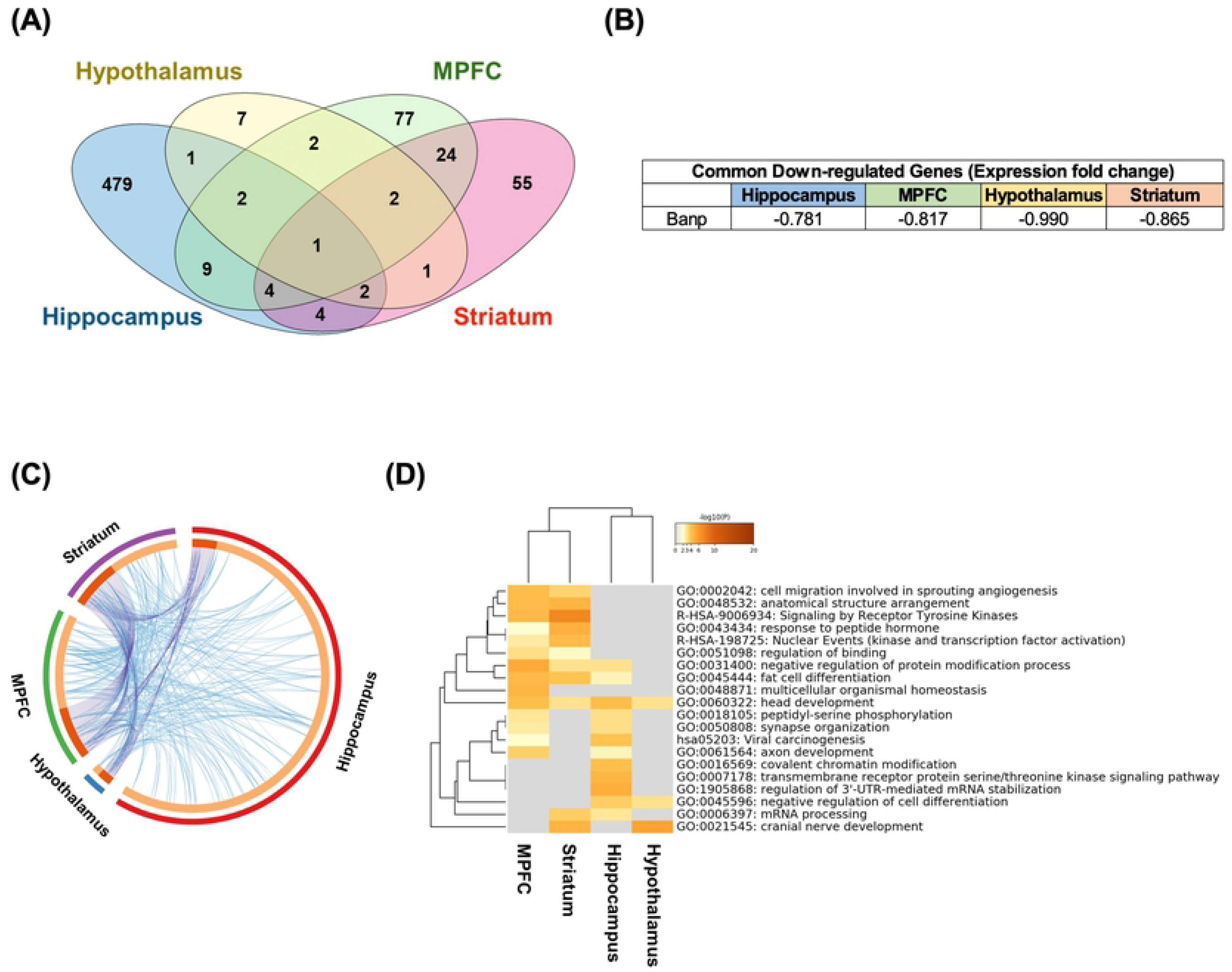
Analysis of the downregulated differently expressed genes for each part of the brain. (A) Venn-diagram for the downregulated differently expressed genes (DEGs) in each part of brain. (B) Commonly downregulated gene in the four parts of the brain and its fold change (log2). (C) Circos plot for the Metascape analysis of the upregulated DEGs. The purple line links the same gene that are shared by multiple gene lists. The blue lines link the different genes where they fall into the same ontology term. (D) Heatmap for gene ontology terms analysis of the upregulated DEGs

For identifying specific gene expression changes induced by sevoflurane inhalation, a comparison was made with the transcriptome array data of the cerebral cortices of sleeping mice as the resembling state[19]. We chose the data of MPFCs exposed to sevoflurane as an equivalent part to the cerebral cortices of the sleeping mice. We extracted DEGs using the same method in our experiments. Regarding the comparison between the gene expression changes in the cerebral cortices of the sleeping mice and those of the waking mice, the sleeping mice had 477 upregulated DEGs and 3572 downregulated DEGs (S5 and S6 Table). As shown in the Venn diagrams, there were 5 common upregulated DEGs and 45 common downregulated DEGs were found between the sevoflurane-anaesthetized and sleeping mice (Fig.4A and B, S7 Table). Moreover, by comparing genes upregulated and downregulated in all parts of the brain exposed to sevoflurane, we found that all the genes except Edn1 were completely expressed differently (Fig.4C and D). These results suggest that most gene expression changes in the brain exposed to sevoflurane might reflect the specific influences of sevoflurane inhalation, and these changes could not be observed in the brains of the sleeping mice.

**Figure4.**
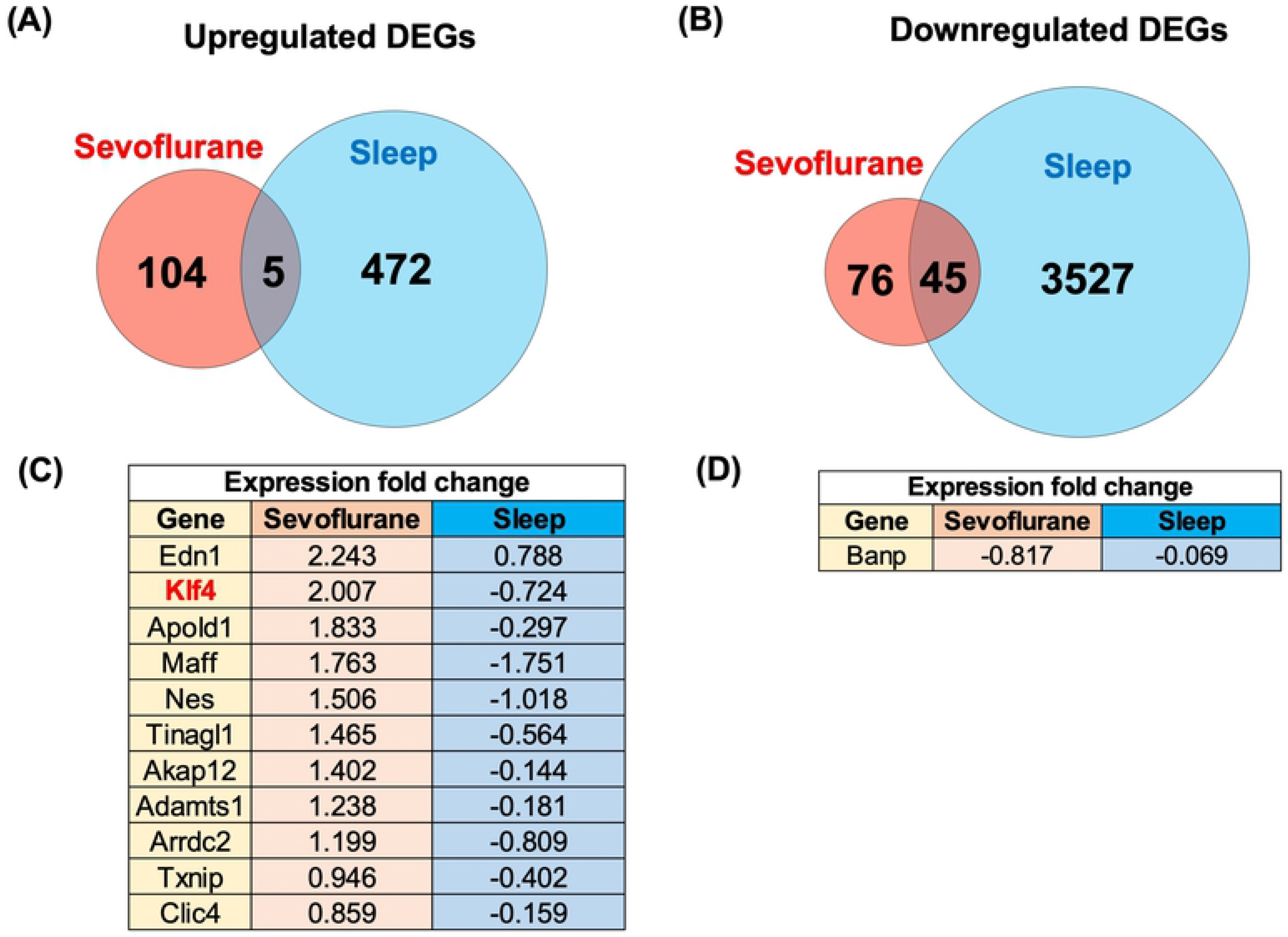
Comparison of differently expressed genes (DEGs) between the brains of the anaesthetized and sleeping mice. (A)(B) DEGs were extracted from the transcriptome array data of the cortical cortices of the sleeping mice. The DEGs in the medial prefrontal cortex of the mice that inhaled sevoflurane and those in the cortical cortices of the sleeping mice were compared. The Venn-diagrams for the upregulated (A) and downregulated DEGs (B) are shown. (C) Table of the expression fold change (log2) of the genes commonly upregulated in the four parts of the brain of the mice that inhaled sevoflurane. (D) Table of expression fold changes (log2) of the genes commonly downregulated in the four parts of brain of the mice that inhaled sevoflurane.

### Identification of KLF4 as a key transcriptional regulator in brain exposed to sevoflurane

As represented by KLF4, sevoflurane induced changes in the expressions of many transcription factors from the analysis of DEGs. Therefore, we hypothesized that sevoflurane changed the activities of specific transcription factors in each part of the brain. To verify this hypothesis, we utilized the wPGSA method [20], with which evaluated the expression changes of the target genes for each transcription factor by using T-scores. A positive T-score means that the transcription factor functions as an activator, while a negative T score means that it functions as a repressor. We regarded transcription factors with both |T-score| > 2.0 and FDR < 0.1 as functional transcription factors. With the wPGSA method, 34, 3, 3, and 1 transcription factors in the MPFC, striatum, hypothalamus, and hippocampus were estimated as activators, respectively. Ninety-three, 188, 113, and 168 transcription factors in the MPFC, striatum, hypothalamus, and hippocampus were estimated as repressors, respectively (Fig. 5A, C, E, and G; S8 Table). Moreover, we identified activators and repressors included in the DEGs, inferring that they particularly functioned owing to the induction by sevoflurane. In the MPFC, the target genes of KLF4, KLF2, and PER2 were upregulated, while those of ATF4 and TAF1 were downregulated (Fig.5B). Likewise, in the striatum, the target genes of 5 transcription factors were downregulated, and in the hypothalamus, the target genes of KLF4 were downregulated (Fig. 5D and F). Finally, in the hippocampus, the target genes of 14 transcription factors were downregulated (Fig.5H).

**Figure5.**
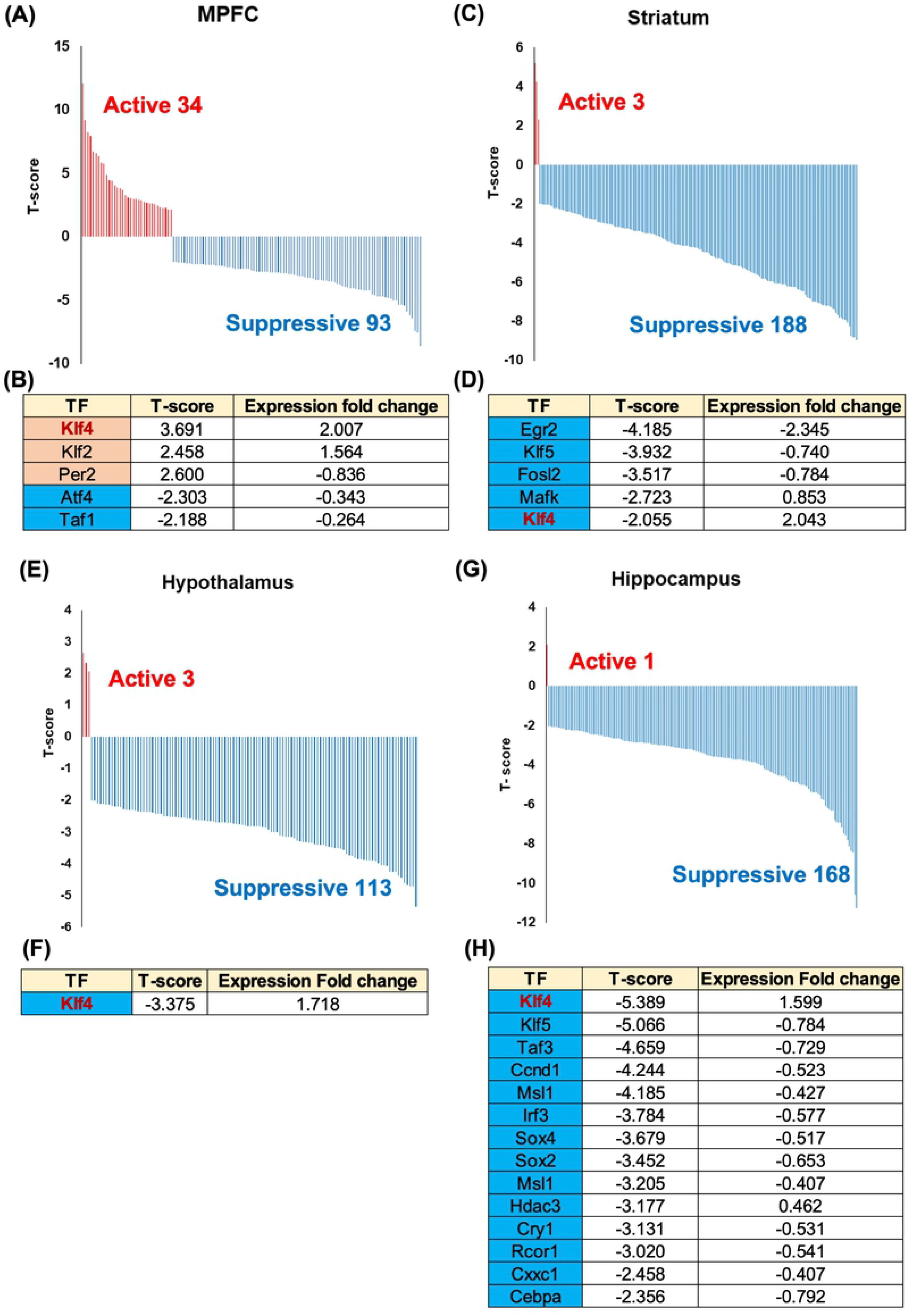
Estimation and comparison of the relative activities of the transcriptional factors. (A)-(H) Weighted parametric gene set analysis (wPGSA) of the fold changes in each part of the brain. Transcription factors (TFs) with T-scores of > 2.0 or < −2.0 were identified and the distributions of the T-scores of the medial prefrontal cortex (MPFC) (A), striatum (C), hypothalamus (E), and hippocampus (G) were drawn. Furthermore, the transcription factors included in differently expressed genes (DEGs) were identified and the tables of the T-scores and expression fold changes for MPFC (B), striatum (D), hypothalamus (F), and hippocampus (H) were made.

Even *Klf4* was upregulated in all four parts of the brain, it worked as an activator in the MPFC, and as a repressor in the other three parts of brain. KLF4 was reported to function as both as an activator and a repressor, and this result might reflect the different transcriptional roles of KLF4 between each part of brain [25]. Moreover, the expression of the same *Klf* family gene, *Klf2*, was also upregulated in the MPFC and functioned as an activator, while *Klf5* and its target genes were downregulated in the striatum and hippocampus. These results indicate the possibility of cooperative functions between the same *Klf* family genes.

Finally, we confirmed the existence of consensus sequences of KLF4 in the DEGs of important functions. The consensus-binding motif sequence of murine KLF4 was GGG(T/C)G(G/T)GGC according to JASPAR (http://jaspar.genereg.net/). On JASPAR, we searched the candidate binding sites of KLF4 in 1000bp upstream sequences for upregulated DEGs annotated GO of “angiogenesis”, and downregulated DEGs annotated GO of “head development”. As shown in the pie charts, 82.7% of the GOs of the upregulated DEGs annotated as “angiogenesis”, and 82.5% of the GOs of the downregulated DEGs annotated as “head development” had consensus-binding motifs in their 1000-bp upstream sequences (Fig. 6A, and B, S9 Table). These results indicate that KLF4 has the potential to regulate the transcription of genes related to angiogenesis and neural development, which might contribute to vascular neogenesis and the appearance of undifferentiated neural cells (Fig. 6C).

**Figure6.**
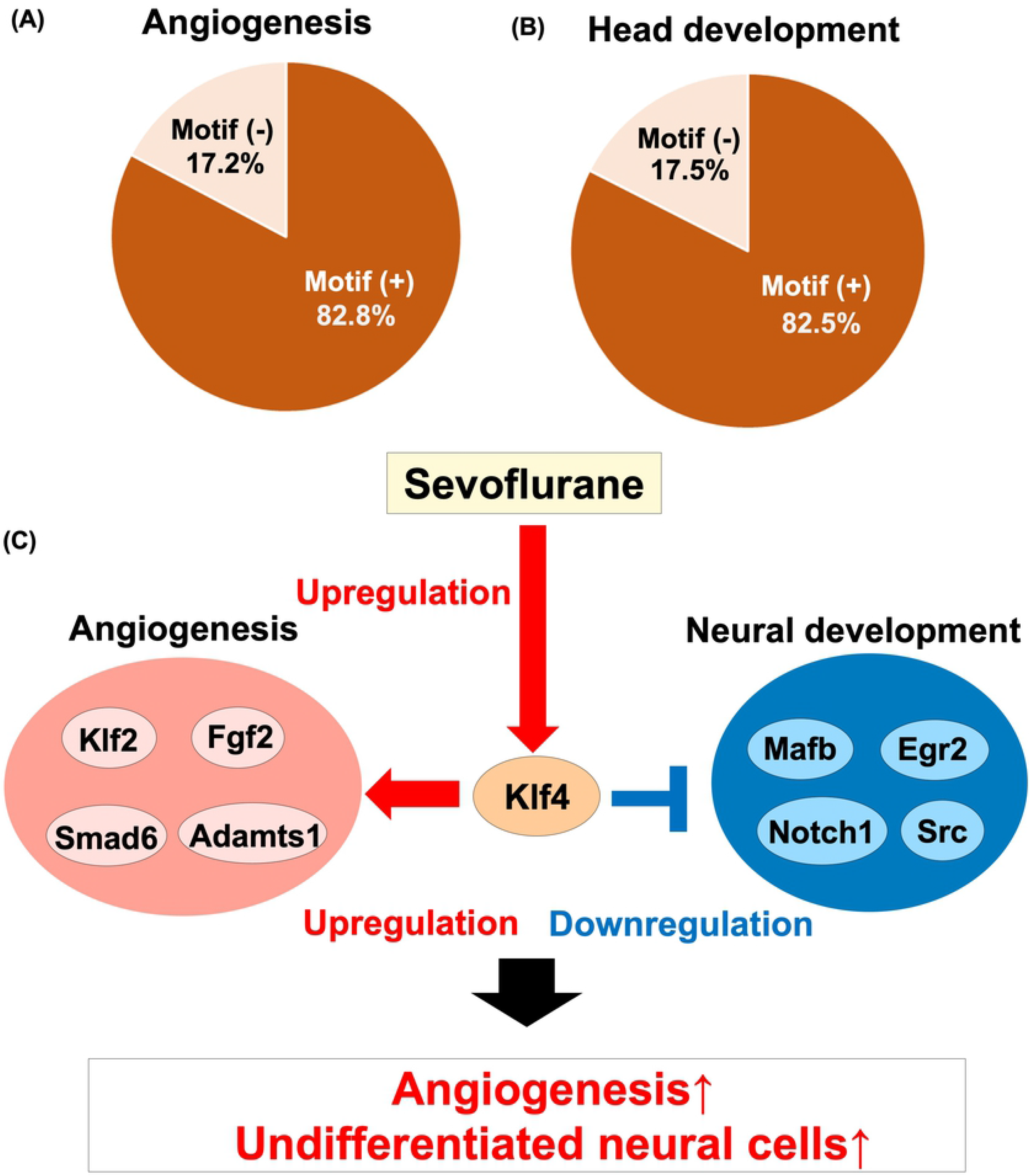
Comparison of the activities of the transcription factors between the brains of the anaesthetized and sleeping mice. (A) The consensus motif sequences of murine KLF4 referred from JASPAR. (B) Pie chart of the existence of KLF4-binding motifs in the 1000-bp upstream sequences of the genes annotated to the GO term “angiogenesis”. (C) Pie chart of the existence of KLF4-binding motifs in the 1000-bp upstream sequences of the genes annotated to the GO term “head development”. (D) Estimated mechanism of the effects of sevoflurane on the brain.

## Discussion

In this study, our group performed a genome-wide transcriptional analysis for the brains of mice that inhaled sevoflurane. Results of our analyses suggest that sevoflurane induced both angiogenesis and the appearance of undifferentiated neural cells in whole brain. These changes in gene expression were not observed in the brains of sleeping mice, and seemed specific to brains exposed to sevoflurane. The transcription factor *Klf4* was commonly upregulated in whole brain, and the results of the wPGSA and motif analysis suggest that KLF4 is a key transcriptional regulator of the angiogenesis and appearance of undifferentiated neural cells.

KLF4 is known as an essential regulator of the initialization of iPS cells, or so-called “Yamanaka factor” [24]. Moreover, the redundant and cooperative functions between KLF2 and KLF5 were reported to be important for sustaining the undifferentiated state of ES cells [26]. Thus, KLF2, KLF4, and KLF5 are known to be fundamental factors for sustaining undifferentiated states. Considering the upregulation of Nestin, which is a specific marker of undifferentiated neural cells, and the decreasing expression of genes associated with neural differentiation, sevoflurane inhalation seemed to cause the appearance of undifferentiated neural cells by the *Klf* family genes.

In the previous report, sevoflurane administration decreased the cerebral blood flow in a statistical parametric mapping analysis [27]. Other reports also indicated that sevoflurane inhalation caused permeability of the brain-blood barrier induced the plasma influx into the brain parenchyma, possibly causing postoperative delirium and cognitive decline [28]. Our results that show the upregulation of genes encoding angiogenesis and the appearance of undifferentiated cells were potentially related with these transient functional changes in the brain caused by sevoflurane. In this context, KLF4 seemed to be the key regulator of these genes, and precise analyses of the roles of KLF4 might be key to unveiling the mechanism of the sevoflurane anaesthesia-induced postoperative functional modification of the brain.

The comparison of gene expressions in the brains of sleeping mice revealed that gene expression changes were specific to the brains exposed to sevoflurane. Especially KLF4 seemed to function specifically by sevoflurane inhalation. The roles of KLF4 seemed to differ among the parts of the brain in our wPGSA. KLF4 has multiple functions, including as activators and repressors, and work context-dependently [25, 29, 30]. Furthermore, our analysis revealed that KLF4 had potentials to upregulate genes related to angiogenesis and downregulate neural differentiation. The variable activity of KLF4 might reflect the differences of these activities between the parts of the brain. For precise understanding of the specific roles of KLF4 induced by sevoflurane, chromatin immunoprecipitation analysis of *Klf4* and histone markers, such as H3K9me3 and H3K27Ac in each part of brain are needed. Nevertheless, our analysis results indicated the importance of KLF4 as a candidate regulator of the effects caused by sevoflurane inhalation.

Our report, which focuses on the changes of transcription factors, provides original and novel approaches for analysing the effects of anaesthetics in brain. This is the first report to evaluate the effects of sevoflurane inhalation, focusing on the activities of transcription factors. As the limitation of this study, we did not evaluate protein expression changes for the genes, and only three of samples were used. Furthermore, we could not exclude the possibility of the effect of the hypoxic condition caused by the respiratory depression induced by sevoflurane [31]. However, our experimental condition (2.5% sevoflurane in 40 % oxygen for 3 hours) is common setting in experiments for studies on the effects of sevoflurane on the brain. None of the genes related to hypoxic reaction, including *Hif1a* and *Arnt,* were detected in our analyses in gene expression changes, supporting the exclusion of the possibility of hypoxia in our experimental condition (S1 Table). For further understanding, proteomic analysis of brains with sevoflurane inhalation and pathological assessment of more samples and oxygen saturation assessment in mice are needed. Nevertheless, our strategies should be better choices for grabbing the whole image of brain activities under an anesthetized condition.

In conclusion, the results of our genome-wide transcriptional analysis of the brains of mice that inhaled sevoflurane suggest the upregulation of angiogenesis and appearance of undifferentiated neural cells in whole brain. Moreover, we identified KLF4 as a potential regulator of the effects induced by sevoflurane inhalation.

## Authors’ contribution

HY designed the study, performed the experiments, analysed the data and wrote the manuscript. YU designed the study, analysed data and wrote the manuscript. TC analysed the data and wrote the manuscript. RK provided critical advices on the data analysis and writing of the manuscript. TM designed the study and provided critical advices on the experiments. CI and HL performed the experiments. TS provided specific skills on preparing the samples. MM analysed data. TU and HA conceptualized the study and were in charge of the overall direction and planning.

## Acknowledgements

We are grateful to all staffs of the Department of Systems BioMedicine at Tokyo Medical and Dental University (TMDU) for their support and discussion.

## Supporting Information

**S1 Fig. RNA-seq analysis for anesthetized brain**

(A)The distribution of log_2_ ((count per million) +4) after normalization.

(B)PCA-plot for RNA-seq data.

**S1 Table. Gene expression data for all the genes in all parts of the brain of mice that inhaled sevoflurane**

Log_2_ (read counts per million +4) of all the genes of all parts of the brain from the RNAseq analysis data by iDEP91 are shown.

**S2 Table. Gene lists of differently expressed genes in each part of the brain**

The gene names and expression fold change data (sevoflurane group vs control group) of the hippocampus, hypothalamus, striatum, and medial prefrontal cortex are shown.

**S3 Table. Lists of genes and gene ontology terms of upregulated differently expressed genes**

Metascape analysis was performed for upregulated differently expressed genes. The gene ontology (GO) terms, their *p* values and genes annotated to each GO terms are shown in the table.

**S4 Table. Genes and gene ontology term lists of downregulated differently expressed genes**

A Metascape analysis was performed for downregulated differently expressed genes. The gene ontology (GO) terms, their *p* values and genes annotated to each GO terms are shown.

**S5 Table. Expression and fold change data for each gene from the transcriptome array data of the cortical cortices of sleeping mice**

The gene names, transcriptome array data and expression fold change data (sleeping group vs control group) from GSE69079 are shown.

**S6 Table. Lists of the differently expressed genes in the cerebral cortices of sleeping mice**

The gene names and each expression fold change data (sleeping group vs control group) for the upregulated and downregulated genes are shown.

**S7 Table. Comparison of the gene expression fold changes of the common differently expressed genes between mice that inhaled sevoflurane and sleeping mice**

The gene names and each expression fold change data for the common upregulated and downregulated genes (sevoflurane group vs control group and sleeping group vs control group) are shown.

**S8 Table. Lists of the transcription factors and their T-scores from the wPGSA for each part of brain**

The activities of the transcription factors (TFs) in the medial prefrontal cortex, striatum, hypothalamus, and hippocampus were calculated using the wPGSA analysis. The T-scores of the transcription factors are shown.

**S9 Table. List of the predicted binding motifs of Klf4 in the upstream sequences of the differently expressed genes**

The predicted binding motifs of *Klf4* for the 1000-bp upstream sequences of the differently expressed genes were identified using JASPAR.

